# Detecting cancer vulnerabilities through gene networks under purifying selection in 4,700 cancer genomes

**DOI:** 10.1101/222687

**Authors:** Anika Gupta, Heiko Horn, Parisa Razaz, April Kim, Michael Lawrence, Gad Getz, Kasper Lage

## Abstract

Large-scale cancer sequencing studies have uncovered dozens of mutations critical to cancer initiation and progression. However, a significant proportion of genes linked to tumor propagation remain hidden, often due to noise in sequencing data confounding low frequency alterations. Further, genes in networks under purifying selection (NPS), or those that are mutated in cancers less frequently than would be expected by chance, may play crucial roles in sustaining cancers but have largely been overlooked. We describe here a statistical framework that identifies genes that have a first order protein interaction network significantly depleted for mutations, to elucidate key genetic contributors to cancers. Not reliant on and thus, unbiased by, the gene of interest’s mutation rate, our approach has identified 685 putative genes linked to cancer development. Comparative analysis indicates statistically significant enrichment of NPS genes in previously validated cancer vulnerability gene sets, while further identifying novel cancer-specific candidate gene targets. As more tumor genomes are sequenced, integrating systems level mutation data through this network approach should become increasingly useful in pinpointing gene targets for cancer diagnosis and treatment.

## 1 Introduction

Over the past decade, the revolution in large-scale cancer sequencing studies has uncovered many driver mutations. These tumor-initiating mutations directly contribute to inducing cancers by activating oncogenes or deactivating tumor suppressors (Garraway and Lander, 2013). Further elucidating key mechanistic, diagnostic, and therapeutic insights into cancer biology, however, could benefit from the identification of genes in networks that are significantly depleted for mutations (those that have undergone purifying selection). Such genes, termed cancer vulnerabilities, are of critical importance to the proliferation, maintenance, and survival of tumors and thus provide an alternative approach to understanding and treating cancer.

Cancer vulnerabilities have traditionally been identified through small-scale experimental studies, often at the single gene level. In these studies, genes were systematically perturbed, and the resulting effect on cancer cell line growth was recorded (Garraway and Lander, 2013; Bass et al., 2009; Lopez and Hanahan, 2002; Etemadmoghadam et al., 2010; Ramsay and Gonda, 2008; Mansouri et al., 1998; Okhrimenko et al., 2005; Kim and Sabatini, 2004). More recently, genome-wide RNAi knockdown and CRIPSR-Cas9 knockout experiments have expanded the study of cancer vulnerabilities in a wide range of cell lines (Cheung et al., 2011; Tsherniak et al., 2017; Wang et al., 2015). This approach has been translated to the clinical stage, by generating a cell line from a clinical tumor sample, investigating this cell line using a combined RNAi and CRISPR-Cas9 approach, and subsequently recommending treatment based on patient-specific vulnerabilities (Hong et al., 2016). However, experimentally generating a comprehensive catalog of vulnerabilities across different cancer cell lines requires considerable resources (e.g. an estimated 600-5,000 samples per tumor type for near-saturation, depending on background mutation frequency) (Lawrence et al., 2014).

Tumor cells accumulate large amounts of passenger mutations as a consequence of unchecked proliferation, genotoxic stress, and defects in the DNA repair machinery. These mutations do not participate in initiating the tumor, and passenger mutation rates can be modeled as randomly distributed across the tumor genome when considering factors such as sequence composition, position, replication timing, transcription-coupled DNA damage repair, and mutation hotspots (Lawrence et al., 2014; Weghorn and Sunyaev, 2017). It should therefore be possible to identify genes that are essential to the proliferation of cancer cells directly from cancer mutation data, by looking for genes that significantly lack passenger mutations.

Several computational methods have accounted for the potential role of passenger mutations in the search for cancer dependencies. Cancer Vulnerabilities Unveiled by Genomic Loss (CYCLOPS) has used available expression data to find genes with partial copy number loss, yielding cells that are vulnerable to their knockouts (Nijhawan et al., 2012). Further, a study on synthetic lethality has analyzed The Cancer Genome Atlas dataset to find mutually exclusive loss-of-function (LoF) gene sets (Ryan et al., 2014). A more recent Bayesian inference approach has successfully found cancer driver genes, but has so far detected only subtle signal from genes under purifying selection in the tumors (Weghorn and Sunyaev, 2017).

Despite these studies’ advancements, the statistical detection of infrequently mutated or preserved genes is strongly confounded by the high background copy number and mutation rates in cancer genomes, potentially unfiltered germline variants in cancer sequencing data, and heterogeneity in mutation rates along the genome, specifically in the context of a single cancer cell. This means that even the most sophisticated methods to detect signals of purifying selection at the gene level are currently underpowered (Tsherniak et al., 2017; Weghorn and Sunyaev, 2017)

In this work, we provide a more robust approach to computationally detect cancer vulnerability genes from cancer genomes, by aggregating weak signals of negative selection across a gene’s first order protein-protein interaction network. We present a statistic, Network Purifying Selection (NPS), which identifies genes that have a network significantly depleted for mutations, indicating that the gene itself is likely to be a vulnerability gene. We applied NPS to 4,742 tumor genomes from 21 tumor types to identify 685 genes with a significant NPS score. Our approach corroborates previously documented studies on cancer dependencies but also identifies a novel set of genes that are likely cancer-specific vulnerabilities. The NPS code is available at www.lagelab.org, and the approach we develop here should become increasingly useful as more cancer genomes are sequenced in the future.

## 2 Results

### Design and properties of the Network Purifying Selection statistic

NPS combines data from 4,742 tumor genomes spanning 21 tumor types and InWeb, a human protein-protein interaction network (Lage et al., 2007; Li et al., 2016), that has been used in dozens of genetic studies, including in the 1000 Genomes Project (Khurana et al., 2013), to calculate the signal of purifying selection in a gene’s functional protein-protein interaction network. Since we specifically wanted a statistic to determine the predictive signal of purifying selection in a gene’s network, we excluded any mutation information on the gene itself in the NPS score calculation. This specific design choice enables NPS to complement any other gene-based method to identify cancer vulnerabilities.

To benchmark the NPS statistic and to test if it accurately classifies cancer vulnerabilities, we defined a set of well established genes from three existing datasets: copy number alterations yielding cancer liabilities owing to partial loss (CYCLOPS), aggregate RNAi gene knockdown data analyzed by DEMETER, and CRISPR-induced LoF essentiality genes from Wang et al. (Figure 1a). Briefly, the CYCLOPS method has identified 56 cancer-specific vulnerability genes by analyzing the effect on tumor growth of knocking out the wild type allele of a gene where the other copy has been lost due to copy number changes in the cancer cells (Nijhawan et al., 2012). DEMETER models gene knockdown effects within the data and computationally subtracts off-target effects to find cancer dependencies (Tsherniak et al., 2017). The CRISPR-induced LoF genes are a set of cell-essential genes required for proliferation and survival in a human cancer cell line (Wang et al., 2015). CYCLOPS, DEMETER, and cancer cell-essential (Wang CRISPR) genes are strongly enriched for the NPS candidates, with odds ratios of 5.4 (P=2.9e-5), 4.7 (P<2.2e-16), and 5.2 (P=2.2e-16), respectively (Supplementary Table 1). To estimate how well the NPS score predicts vulnerability genes and to see if genes that do not pass the significance cutoff might still be valuable candidates, we also calculated receiver operating characteristic (ROC) curves for all three sets. AUCs were 0.68 (P=0.153), 0.677, and 0.645 (P=0.0139), respectively (Figure 1b).

**Figure 1.**
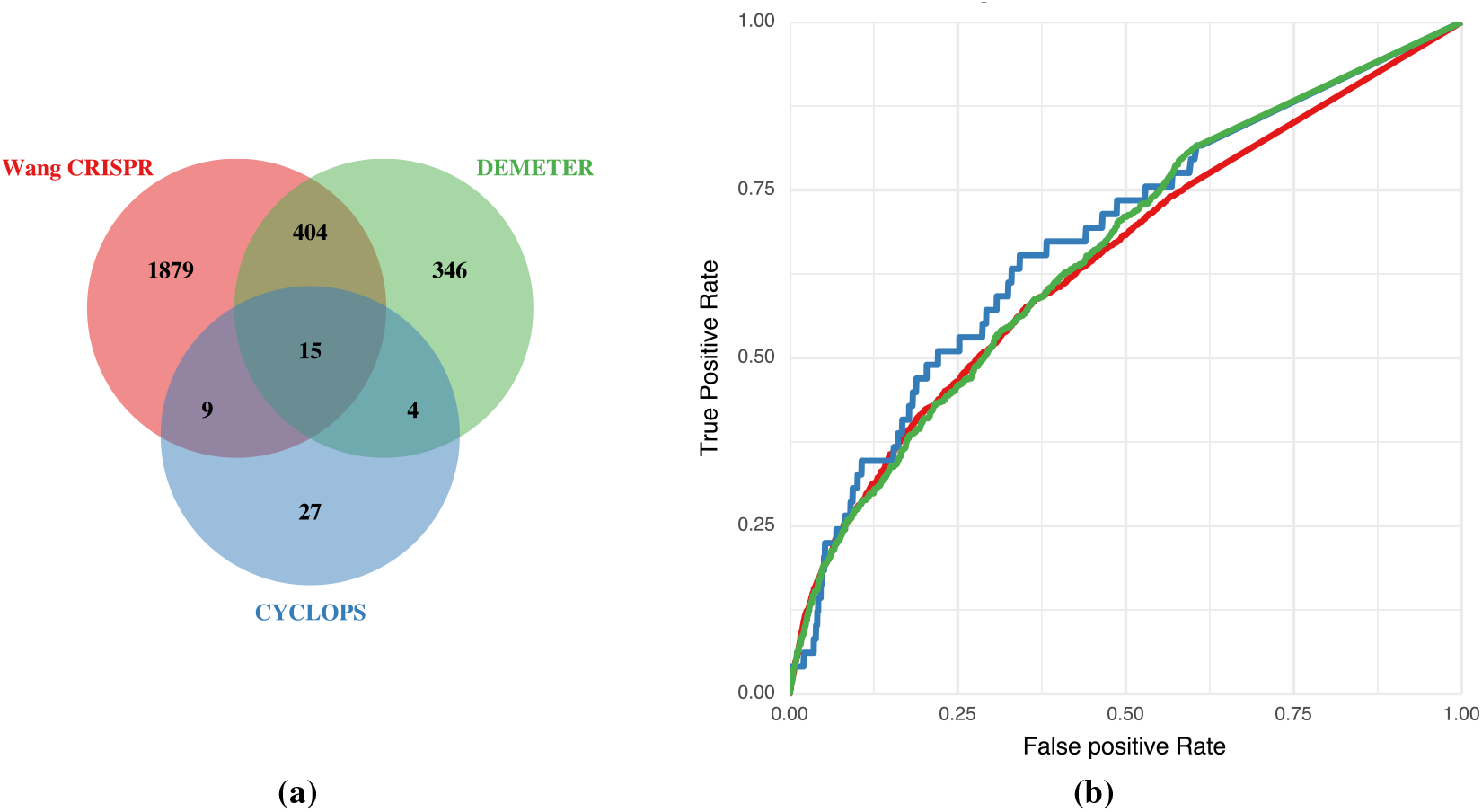
NPS distinguishes true cancer vulnerability genes. **(a)** Overlap of the three cancer benchmarking gene sets. **(b)** Receiver operating characteristics (ROC) curves for genes in CYCLOPS (blue), DEMETER (green), and cancer cell-essential genes (red). The areas under the ROC curves (AUCs) are 0.68, 0.677, and 0.645, respectively. Genes from the CYCLOPS, DEMETER, and CRISPR-induced LoF cell essentiality cancer sets are defined as true positives, and all other genes in InWeb as true negatives. NPS scores are calculated for all genes using pan-cancer data and used as a classifier.

The majority of genes scored by NPS fit the null hypothesis and lie on the diagonal on a quantile-quantile plot, confirming that the NPS accurately calculates the significance level of purifying selection in the neighborhood of an index gene (Supplementary Figure 1). To correct for knowledge contamination (where well known genes are better studied, leading to increased interaction data), we apply a permutation scheme that considers the network topology of the neighborhood to account for underlying confounders when permuting the neighboring genes. This increases the signal to noise ratio and demonstrates that NPS adequately normalizes for the number of interactions that a gene has at the protein level (Supplementary Figure 2).

These findings indicate that genes significant in the NPS are enriched for previously associated cancer genes. The NPS score thus accurately distinguishes CYCLOPS, DEMETER, and cancer cell-essential genes from other genes covered by interactions in the InWeb database.

### Predicting Network Purifying Selection candidates from tumor genomes

To test if the NPS statistic can predict new vulnerability genes from existing cancer genome data, we calculated NPS scores for all genes that had at least one high-confidence protein interaction in InWeb. We declared genes as significant at a false discovery rate of *Q* ≤ 0.1 using the pan-cancer cohort of 4,742 tumors. The pooled set (named NPS5000, Supplementary Table 2) contains all unique genes that were significant in the pan-cancer analysis or in at least one of the 21 tumor types. NPS5000 is comprised of 685 genes, many of which are linked to known cancer biology.

Using the Molecular Signatures Database of annotated gene sets (Liberzon et al., 2011), one hundred and ninety one canonical pathways were identified as being enriched (FDR, q≤0.1) for the NPS500 set. The functional roles of these genes are primarily in the following categories: transcription & translation, cell cycle, cell signaling, metabolism, immune response, and membrane & transport (Supplementary Figures 3 and 4, Supplementary Table 3). As expected, many of these processes are found in subsets of living cells, but also in the hallmarks of cancer.

### Network Purifying Selection detects cancer-specific vulnerabilities

NPS should in theory identify both genes that specifically are cancer vulnerabilities, but also genes that are generally under purifying selection in human populations, because they are essential to both normal and cancer cells. To dissect this phenomenon, we tested the overlap of our NPS5000 set with genes from population genetic studies that are known to have a high probability of being intolerant to LoF mutations (Samocha et al., 2014) (Figure 2). Specifically, we overlapped the NPS5000 set with genes from the Exome Aggregation Consortium (ExAC) population exome database that have a probability of intolerance to LoF (defined as pLI) ≥ 0.9 (Lek et al., 2016). ExAC contains the exomes of 60,706 people, loosely representing the general population, and lets us estimate gene tolerance to LoF mutations at a population level.

**Figure 2.**
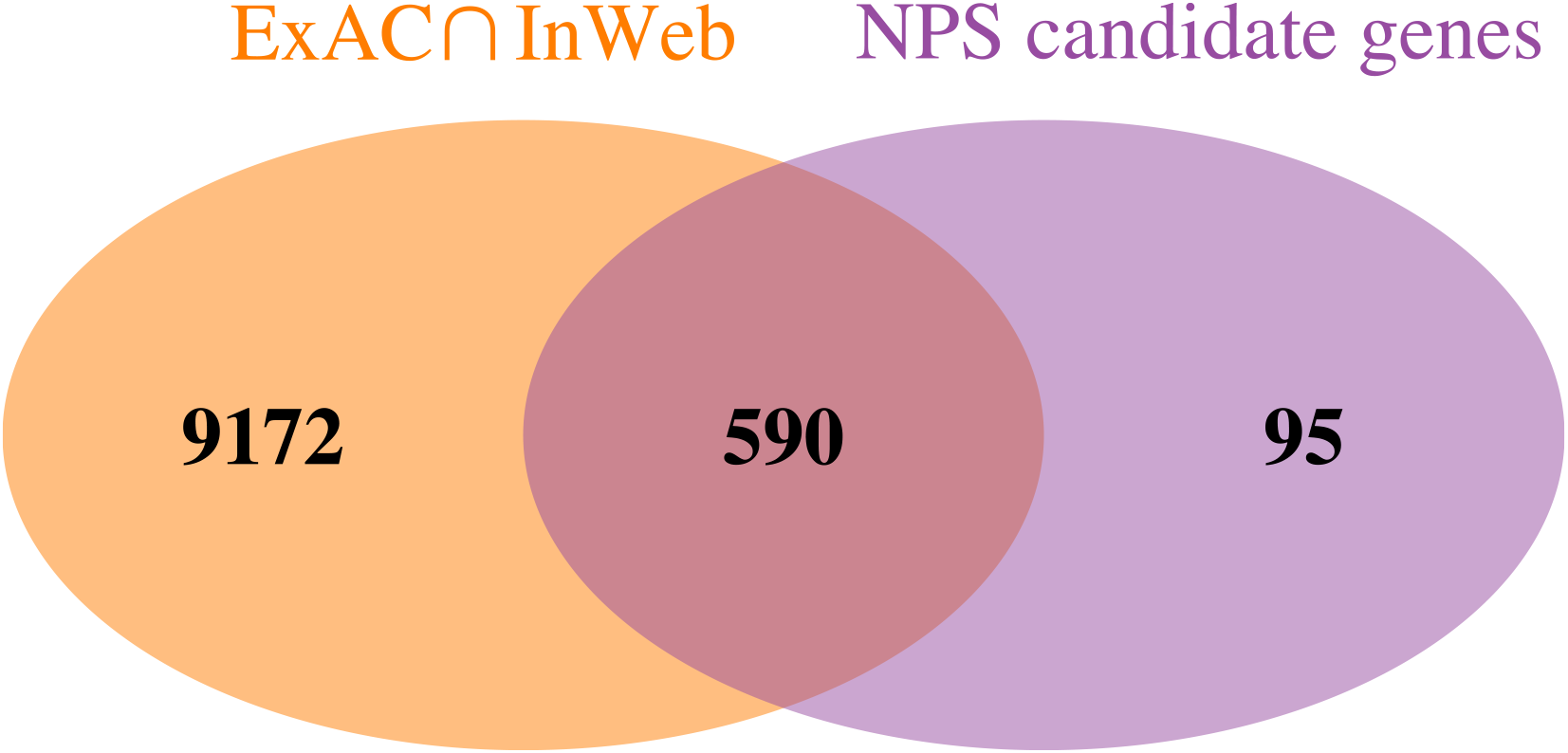
NPS identifies both disease state-agnostic (orange) and cancer-specific (purple) vulnerabilities. ExAC genes included are those with pLI scores > 0.9.

After filtering ExAC (16,453 genes) for genes present in InWeb3 (12,000 genes), 590 of the 685 NPS-imputed genes were linked to ExAC vulnerabilities in response to LoF alterations. The remaining 95 genes were found to have no significant LoF consequences in the general population (Supplementary Table 2).

### Tumor essentiality of NPS candidates

We assessed the impact of knockouts of three gene sets on the growth of 216 cancer cell lines. These sets comprise all ExAC genes (Background), the overlapping NPS and ExAC vulnerability genes, and the non-ExAC NPS vulnerability genes (e.g. those genes specific to cancer dependencies) (Figure 3). We used the cell proliferation data compiled through Project Achilles, which catalogues the genetic vulnerabilities of genomically characterized cancer cell lines through individual gene knockouts using the CRISPR-Cas9 system (Cowley et al., 2014); internal Broad data set).

**Figure 3.**
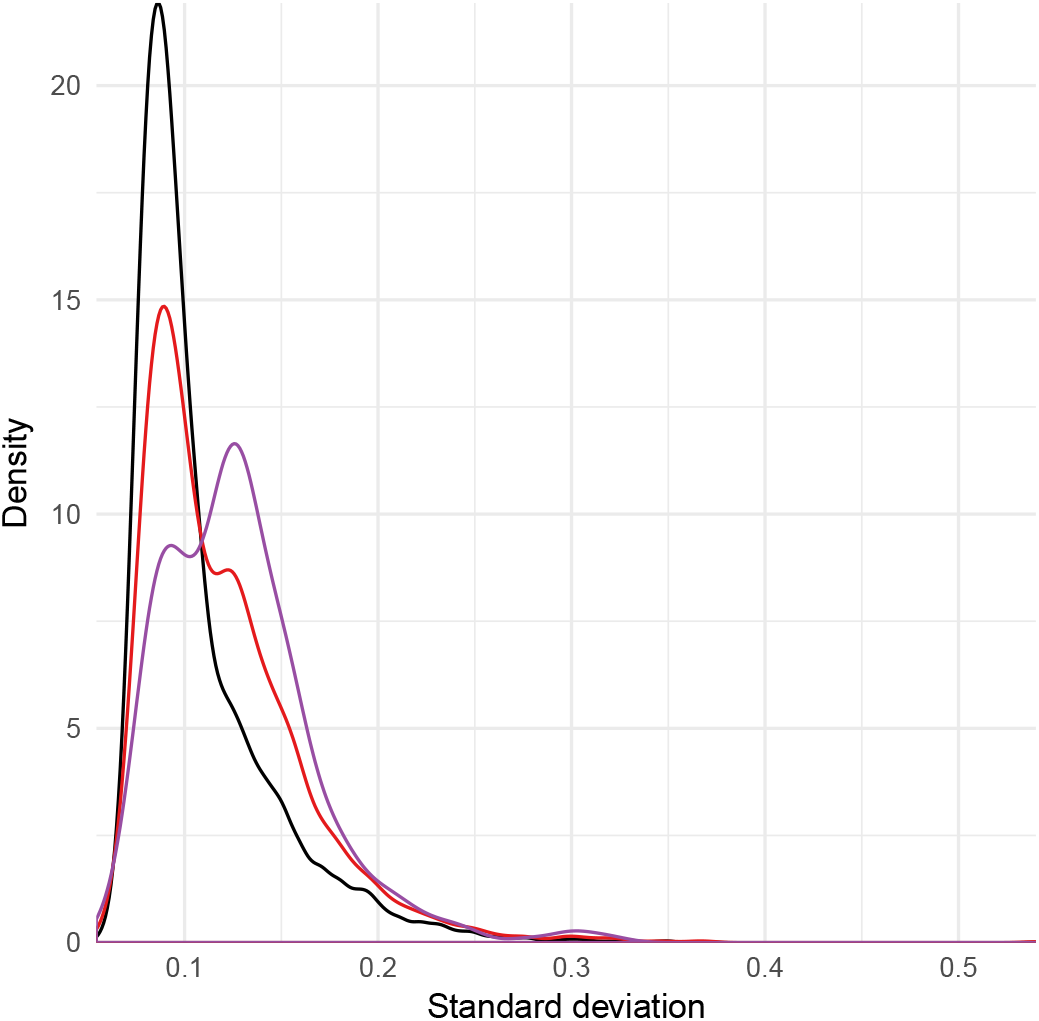
Tumor growth divergences resulting from gene knockouts. Aberrant tumor growth resulting from the loss of function of the aggregate NPS-imputed + ExAC vulnerability genes (red), and non-ExAC NPS vulnerability genes (purple), as it diverges from the null distribution (aggregate ExAC genes) of tumor growth (black).

The aggregate set of genes in cancer networks under purifying selection impacts tumor proliferation at a level that is statistically significantly different from that of background gene candidates. This difference is non-directional (knockout of a gene enhances or decreases proliferation); therefore, we calculated the standard deviation from the normal tumor growth rate as an absolute measurement. The Background set results in a Gaussian distribution, ExAC+NPS gene knockouts lead to a slight deviation from normal growth, and the NPS-imputed, non-ExAC gene knockouts (cancer-specific vulnerabilities) yield the most starkly aberrant tumor growth (Figure 3). Overall, knocking out genes from the set of 685 leads to aberrant tumor cell growth and statistically varied tumor proliferation.

## 3 Discussion

The NPS-imputed gene statistic described here facilitates the identification of novel, previously hidden cancer vulnerabilities and broadens the pool of potential drug targets. Due to the network-based approach to calculate the NPS statistic, those genes identified as cancer dependencies have NPS scores that are not influenced by the gene itself, but rather, by the mutation landscape of its direct protein interaction network. This approach eliminates the possibility of any bias from the alteration rate of an index gene and enables NPS to complement previously established cancer vulnerability detection methods.

Benchmarking analyses comparing the NPS-imputed genes with CYCLOPs, DEMETER, and CRISPR-induced cancer cell-essential genes reveal significant enrichment in all three of these selection lists and significant ROC AUCs, indicating that a substantial subset of the genes identified through this network approach are known cancer dependencies.

The overlap between NPS-imputed genes and those in the ExAC database indicates that the NPS method has identified genes crucial to proper cell functioning, for which a high preservation is essential. This overlap analysis suggests that LOF alterations in the 95 genes only significant in NPS and not in ExAC may be linked to tumor-specific dependencies.

Statistically significant tumor growth aberrations due to NPS gene knockouts, as determined by Achilles gene knockout studies (Meyers et al., 2017), indicate the direct relevance of these genes in sustaining tumor proliferation. Knockouts of NPS genes change tumor growth rate, and NPS only, non-ExAC gene knockouts (those that are putative cancer-specific vulnerabilities) affect tumor growth even more, suggesting the identified genes’ key role in tumor development.

The pan-cancer NPS-imputed gene analysis presented here unveils that genes in networks under purifying selection in cancers could be harnessed for cancer treatment. The existence of FDA-approved drugs targeting these newly identified cancer vulnerabilities (Supplementary Table 4) supports the claim for repurposing therapies, enabling a shortened timeline to treat otherwise intractable cancers.

## 4 Methods

### Calculating the network purifying selection score

For a given index gene, the Network Purifying Selection statistic is formalized into a probabilistic score that reflects the index-gene-specific composite purifying selection (i.e. the aggregate of single-gene MutSig suite Q values from Lawrence et al.) across its first order biological network and is calculated via a three-step process. First, we identify all genes it interacts with directly at the level of proteins, only including high-confidence quality-controlled data from the functional human network InWeb (where the vast majority of connections stem from direct physical interaction experiments at the level of proteins). Second, the composite purifying selection score across members of the resulting network is quantified by aggregating single-gene MutSig suite Q values from Lawrence et al. into one value ϕ using an approach inspired by Fisher’s method for combining p-values:

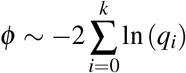

Where q_i_ is the MutSig suite Q value for gene i, and k is the amount of genes in the first order network of the index gene (i.e. the index gene’s degree). Third, by permuting the InWeb network using a node permutation scheme, we compare the aggregated burden of mutations θ to a random expectation. In this step, the degree of the index gene, as well as the degrees of all genes in the index gene’s network is taken into careful consideration. The final NPS score of an index gene is therefore an empirical P value that reflects the probability of observing a particular composite mutation burden across its first order physical interaction partners (at the level of proteins) normalized for the degree of the index gene as well as the degrees of all of its first order interaction partners. Because we are interested in estimating the purifying selection independent of the index gene, this gene is not included in the analysis and it does not affect the NPS calculation.

### Classifying cancer vulnerability genes

For each gene represented in InWeb (12,507 or 67% of the estimated genes in the genome), we used the gene-specific NPS probability to classify it as a cancer vulnerability gene or not. True positive genes were a set of CYCLOPS, DEMETER, and CRISPR-induced cell-essential cancer genes. Specifically, we selected 56 genes categorized as CYCLOPs genes as described in (Nijhawan et al., 2012), and DEMETER genes were defined as the 769 genes that were differentially required in subsets of the analyzed cell lines at a threshold of six SDs from the mean (Tsherniak et al., 2017). True negatives were defined as all genes in InWeb that were not in these three sets, which is likely conservative, as we currently lack the power to confidentially estimate the number of potential vulnerabilities. We used the NPS probability as the classifier and calculated the AUC for each gene set.

### Testing the robustness of the NPS approach

A more in depth evaluation has been performed in Horn et al. Here, to assure that the permutation holds up, we limited testing to compare permutation methods (random permutation vs. connectivity aware). We ran the full analysis using both approaches and compared the quantile-quantile plots. This analysis confirmed that ignoring the interaction structure of the immediate neighborhood significantly depletes the signal (Supplementary Figure 1).

### Generating the NPS5000 set

NPS probabilities were determined for every gene in InWeb that was covered by interaction data using 10^6^ permutations. The FDR Q values were calculated as described by Benjamini and Hochberg, based on the nominal P values controlled for 12,507 hypotheses. We performed NPS analyses with the pan-cancer Q values, as well as Q values from each of the 21 tumor types for which they were available. As it is a technical limitation of the NPS approach that it is currently not possible to make 5.5 × 10^6^ network permutations, we could not create a dataset where we correct for all 12,500 × 22 hypotheses tested in the NPS5000 set.

### Detecting cancer-specific vulnerabilities

To expand the known catalog of vulnerabilities, we tested whether our approach could predict both putative and novel, cancer-specific vulnerability genes directly from tumor and exome sequencing data (ExAC). We filtered for ExAC genes with pLI scores ≥ 0.9 and checked for their overlap with NPS genes, as well as for NPS genes that did not fall in the highly potent ExAC gene LOF list, for cancer-specific targets.

### Dissecting tumor gene essentiality

We tested the effects of NPS gene knock-outs on cancer cell lines using the Project Achilles cancer cell line gene knockout database. For the following groups of genes, we assessed the change in growth rate from base-line across 216 cancer cell lines when those genes were knocked out: the 590 NPS + ExAC genes, the 95 NPS only (non-ExAC) genes, and all ExAC genes. For each of these three gene knockout groups, we calculated the resulting standard deviations of growth from the null distribution of cancer cell proliferation rates.

## 5 Acknowledgments

We would like to thank Professor Aviv Regev for her inputs on the work.

